# Age-dependent shift from high lipid content to high haemolymph sugar levels in *Bombus terrestris* (Hymenoptera: Apidae)

**DOI:** 10.64898/2026.01.17.700087

**Authors:** Helena Schulte, Carolin Bäuml, Erhard Strohm, Christoph Kurze

## Abstract

Division of labour is key to the success of eusocial insects. While honeybees are the primary example of age-based polyethism, the physiological changes that may contribute to task allocation in bumblebees remain poorly understood. To characterise potential age-related changes energy stores, we quantified the haemolymph sugar concentrations (fructose, glucose and trehalose) and body lipid content in *Bombus terrestris* workers of five age classes (0-28 days) and varying body sizes (head width, intertegular distance and dry mass, which were strongly correlated). We found a pronounced transition during the first week of adult life. Newly emerged workers (callows, ≤ 1 day) had high absolute and relative lipid content (% of dry mass) but low haemolymph sugar concentrations, whereas workers aged 7 days or older had about half the body lipid content and approximately 6-fold higher glucose and 4-fold higher trehalose concentrations. Fructose did not differ between age classes. Body size (head width) was positively associated with absolute lipid content in callows and, less robustly, with trehalose concentration, but age explained most of the variation. Our findings indicate an early, age-related shift in energy stores in bumblebee workers and highlight the importance of accounting for worker age in social insect research.

## 1. Introduction

Key to the success of eusocial insects (i.e. ants, termites, social wasps, and social bees) is the division of labour, or task allocation [1-3]. While this typically is characterised by a clear division between reproductive (queens, males) and non-reproductive tasks (workers) such as brood care and foraging, our understanding of task allocation in the primitively eusocial bumblebees remains limited [4-7]. Workers of *Bombus terrestris* (Hymenoptera: Apidae) have been shown to also lay unfertilised, haploid eggs that develop into males (arrhenotokous parthenogenesis) [8, 9], and worker mating has been reported by one study [10]. Resource allocation is expected to be crucial for successful worker reproduction, yet little is known about how energy reserves such as lipid stores (important for ovary development [11]) and haemolymph sugar titres (relevant for flight [12, 13]) change as workers age.

In the honeybee *Apis mellifera* (Hymenoptera: Apidae), workers transition from in-hive tasks to riskier tasks outside the nest, such as foraging, as they age [3]. This transition, known as age-based polyethism, is mediated by physiological changes such as increasing juvenile hormone (JH) titres [14, 15], accompanied by a shift from high lipid reserves [16] to increased mobilisation of trehalose [17]. Trehalose is the major circulating energy source essential for fuelling honeybee flight and is a useful indicator of nutritional status [18, 19].

In contrast, the importance of age for task allocation in bumblebee workers is less clear [4, 20-22]. Task allocation in bumblebees has been found to be associated with body size (alloethism), with smaller workers tending to remain inside the nest and larger workers to do riskier tasks outside [4, 5, 20]. This aligns well with the general body size/foraging range hypothesis, where larger body size enables bees to forage over longer distances [23, 24]. While larger body size may increase flight speed in *B. terrestris*, age may more strongly influence flight distance than body size as demonstrated in laboratory experiments using flight mills [7]. Flight is not only energetically costly [13], but increased foraging activity has been reported to lead to reduce longevity in honeybees [25-27] and bumblebees [4]. Body size is also expected to influence energy metabolism, because metabolic rate and energy storage capacity scale with body mass, and body size explains approximately half of the variation in flight metabolic rate in *Bombus impatiens* workers [28]. In addition, it has been proposed that large bumblebee workers may maintain the potential to reproduce by avoiding these risky and energy-intensive tasks, thereby maintaining or build up higher lipid reserves, which may support the development of larger oocytes [6].

Despite energy reserves playing an essential role in flight performance and potentially also for task allocation in bumblebees, it remains unclear how haemolymph sugar levels and lipid stores are affected by worker age and body size. Here, we addressed this gap and hypothesised that: (1) trehalose levels in the haemolymph increase as workers age, which may be relevant for flight capacity (i.e. increased fuel availability for longer flights); and (2) lipid reserves decrease with age, but large individuals may maintain higher lipid reserves, which have been proposed to be needed for ovary development [6]. We only focused only on age- and size-related physiological patterns that may be relevant for task allocation, and did not observe behaviour, assess ovary development or measure hormone titres to test task allocation directly.

## 2. Materials and Methods

### 2.1 Bumblebee husbandry

We maintained six commercial queen-right *Bombus terrestris* colonies (Natupol Research Hives, Koppert B.V., Netherlands) under standard laboratory conditions (23 ± 1°C, ~ 40% RH, and 14:10 h light:dark rhythm) [7]. Each colony was housed in a nest box (length x width x height: 27 x 24 x 14 cm), which was connected to a small foraging arena (60 x 40 x 28 cm), allowing aging bees to fly. These bees had *ad libitum* access to 40% w/v sucrose solution. Additionally, each colony was supplied with 6-11 g of pollen candy (67% w/w organic pollen, 25% w/w sucrose, 8% w/w water) daily.

### 2.2 Ethical statement

This study was conducted in accordance with the ethical principles of the German Animal Welfare Act (Tierschutzgesetz, TierSchG). Formal institutional ethical approval was not required for experiments with insects (§ 7 to § 9 TierSchG). All procedures followed the principle of minimising pain, suffering, or harm without scientific justification (§ 1 TierSchG).

### 2.3 Marking

Newly emerged workers (≤ 1 day post-emergence; hereafter callows) were collected and marked using water-based paint markers (5M Uni-Posca, Mitsubishi Pencil France S.A., France) similar to previous research [7], allowing us to identify bees from specific cohorts at sampling. Individual sex was confirmed by counting antennal segments under a stereo microscope (workers have 12, males 13). Subsequently, workers were returned to their natal colony until aged 7, 14, 21, or 28 days. Individuals assigned to the 0-day-old age class (≤ 1 day, callows) were used directly.

### 2.4 Sampling

We measured haemolymph sugars, head width, dry mass and lipid content from each individual bee. In total we collected 159 marked bumblebee workers aged 0 (≤ 1 day, callows), 7, 14, 21, and 28 days (n = 32; 31; 32; 33; 31) from four to six colonies (Col#1-6: n = 8; 21; 27; 47; 34; 22; four to six colonies per age class) (supplementary Table S1). To account for this sampling imbalance across colonies, we included colony ID and collection date as random intercepts in all mixed-effects models (see statistical analyses) and conducted a sensitivity analysis by refitting each final model after removing one of the six colonies at a time (supplementary Table S2).

Prior to sampling their haemolymph, we kept bees individually in 20 mL glass vials, closed with a mesh lid for three hours to allow each bee to feed *ad libitum* on 40% sucrose solution. Each vial contained a cardboard floor (4 x 1.5 cm). Afterwards, we anaesthetised each bee with CO_2_ for about 10 s and carefully punctured it with a disinfected needle in the intersegmental skin between the third and fourth sternite (ventral abdominal segments) to collect haemolymph with a 1 μL capillary (Microcaps^®^, Drummond Scientific Company, USA). The haemolymph samples were immediately diluted in 99 μL Millipore water (1:100 dilution) and stored at −75°C until GC-MS analysis. The bumblebees were directly frozen at −20°C in individual 2 mL reaction tubes until morphometric measurements.

### 2.5 Preparation of sugar standards and haemolymph samples

Prior to GC-MS (gas chromatography-mass spectrometry) analysis, haemolymph samples and sugar standards (for glucose and sucrose from 0.01 to 1 mg/mL; fructose and trehalose from 0.01 to 1.5 mg/mL) were derivatised following previously described protocols with slight modifications [29, 30]. First, 30 μL of each sample was transferred to a new 1.5 mL reaction tube (Eppendorf GmbH, Germany), lyophilised for 4 hours (Lyovac GT2, Leybold-Heraeus GmbH, Germany), and immediately stored at −75°C for up to 3 days. Then 30 μL of 0.05% hexachlorobenzene pyridine solution (0.5 μg/μL; Sigma-Aldrich Chemie GmbH, Germany) was added to each dried haemolymph sample as an internal standard (IS). To prevent the formation of multiple derivates during silylation (derivatisation), reducing sugars in samples were first oximated by adding 10 μL of 5% w/v methoxyamine-hydrochloride in pyridine (Sigma-Aldrich Chemie GmbH) and incubating samples for 30 min at 75°C. For silylation, 10 μL of BSTFA with 1 % TMCS (N,O-Bis-(trimethylsilyl)-trifluoroacetamide with 1% trimethylchlorosilane; Sigma-Aldrich Chemie GmbH) was added to each sample and incubated for another 30 min at 75°C. Samples were stored at −75°C for up to 3 days until GC-MS analysis.

### 2.6 GC-MS Analysis

Prior to GC-MS analysis, each sample was diluted with 340 μL of DCM (dichloromethane; Fisher Scientific GmbH, Germany) and 190 μL of the solution was transferred to a GC-MS glass vial and directly placed into an autosampler (Agilent 7683, Agilent Technologies Inc., USA). Samples and sugar standards were analysed with a gas chromatograph (Agilent 6890N) connected to a mass selective detector (Agilent 5973). The injector temperature was maintained at 300°C and the mass spectrometry source temperature was maintained at 230°C. The chemical compounds in 1 μL of sample were separated on a HP-5MS capillary column (length = 30 m, diameter = 0.25 mm, film = 0.25 μm, Agilent Technologies Inc.) with helium as carrier gas at a constant flow of 1 mL min^-1^. The GC oven had an initial temperature of 60°C held for 1 min and was ramped at 5°C min^-1^ until reaching 300°C, which was held for 10 min. Electron ionisation mass spectra were recorded at an ionisation voltage of 70 eV.

The resulting chromatogram for each sample was analysed using manual peak integration (Agilent MSD Productivity ChemStation). The masses and retention times for the main sugar peaks are provided in supplementary Table S3. Calibration curves were produced by calculating the ratio of the integration area of the sugar standard over the integration area of the IS across all concentrations and fitting a linear regression line forced through the origin (supplementary figure S2). Sample concentrations were calculated by dividing the sample sugar to IS peak area ratio by the calibration slope. To obtain the original sugar concentrations within each sample, concentrations were multiplied by 100, accounting for the 1:100 dilution.

### 2.7 Measuring body size proxies and lipid content

Body size proxies (head width, intertegular distance = ITD, and dry mass) and whole body lipid content were determined for each bee following our previous protocols [31, 32]. The head width (maximal distance between the lateral margins of the compound eyes in frontal view) and ITD (distance between the tegulae) of each bee were measured using a digital microscope (VHX-500F, Keyence GmbH, Germany). Afterwards, whole-body dry mass and lipid content were measured on the same specimen. To facilitate drying and solvent penetration, we first cut each bee from the stinger to the fourth sternite. The initial dry mass (including body lipids) was determined by drying the specimens at 60°C for three days and weighing them (d = 0.1 mg, M-Pact AX224, Sartorius GmbH, Germany). To determine the lipid content of the same specimens, the entire body lipids were extracted in 5 mL petroleum ether for five days, the extracts were discarded, the specimens were rinsed with fresh ether, dried again for another three days at 60°C and re-weighed (i.e. dry mass without lipids). The absolute lipid content was calculated as the difference between the initial dry mass (including body lipids) and the dry mass after lipid extraction. The relative lipid content is the absolute lipid content relative to the initial dry mass (in %).

### 2.8 Statistical analyses

All statistical analyses and data visualisations were performed using R version 4.5.1 [33]. The dataset and R Markdown code and output used for statistical analyses and figures, including all model details, validations and pairwise comparisons, are available via Zenodo (https://doi.org/10.5281/zenodo.18282075).

The relationships between the body size proxies ‘head width’ and ‘dry mass’ and between ‘head width’ and ‘ITD’, and potential differences in body size among ‘age’ classes, were assessed by fitting generalized linear mixed-effects models (GLMMs) with a Gaussian distribution using the *glmmTMB* package [34] and by Pearson’s correlations between body size proxies.

The effects of ‘age’ and ‘head width’ as the selected body size proxy (highly correlated with ‘ITD’ and ‘dry mass’, see figure 1) and their interaction on haemolymph sugar concentrations (i.e. ‘fructose’, ‘glucose’, ‘trehalose’) and body lipid content (i.e. ‘absolute lipid’ and ‘relative lipid’ content) were tested by initially fitting GLMMs with Gaussian distribution. Those GLMMs included ‘colony ID’ and ‘collection date’ as random intercepts to account for the non-independence of workers from the same colony, collection date, and for the imbalanced sampling across colonies. Furthermore, we conducted a sensitivity analysis by refitting each final model after removing one of the six colonies at a time to test the robustness of our observed effects, using a planned contrast between callows and the mean of all older age classes (supplementary Table S2).

**Figure 1.**
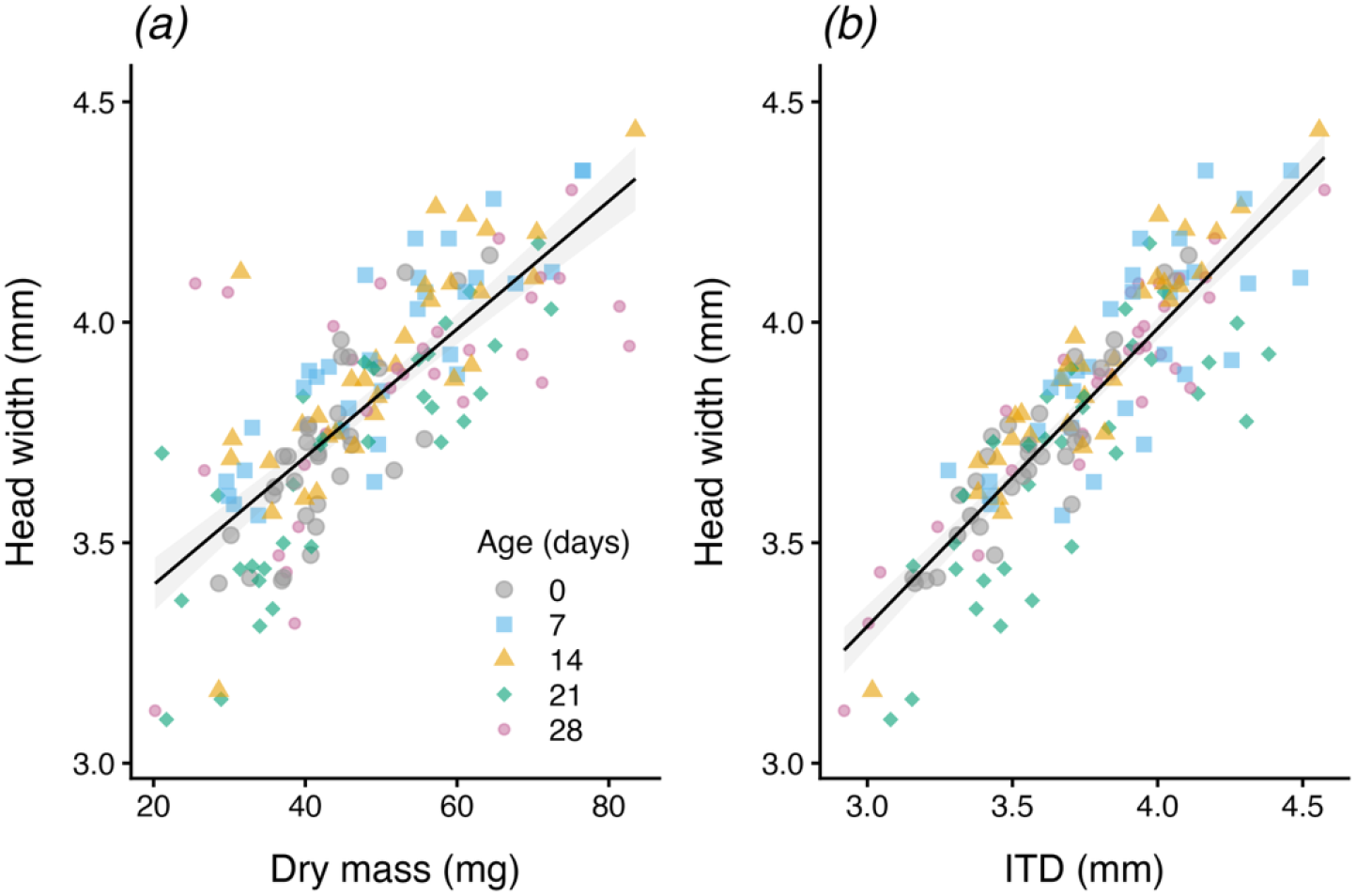
Relationships between body size proxies in bumblebee workers across five age classes. Scatter plots confirm the strong positive relationship between body size proxies: dry mass (mg), head width (mm), and intertegular distance (ITD in mm). ***(a)*** Head width (mm) increased with dry mass (mg) (Pearson’s *r* = 0.77, *t* = 14.95, *df* = 157, *p* < 0.001). ***(b)*** Head width (mm) increased with ITD (mm) (*r* = 0.87, *t* = 22.58, *df* = 157, *p* < 0.001). Lines show linear regressions across all workers (*n* = 159 from 6 colonies, see also see supplementary figure S1 and Table S1) with 95% confidence intervals. Age classes are denoted by distinct symbols and colours: 0 d (callows; grey circles, *n* = 32), 7 d (sky blue squares, *n* = 31), 14 d (orange triangles, *n* = 32), 21 d (bluish-green diamonds, *n* = 33), and 28 d (reddish-purple small circles, *n* = 31).

Model selection was based on the Akaike information criterion (AIC), with lower AIC values indicating the most parsimonious and best-fitting model. The best-fitting models were compared to a simpler nested model using likelihood ratio tests (LRTs). When added terms did not improve model fit significantly, the simpler model was chosen and compared against its respective null model using LRT. Model assumptions and dispersion of the data were checked using the *DHARMa* package [35]. The ‘glucose’ model was fitted with a Gamma (log-link) distribution and ‘age’ as a dispersion term to meet model assumptions. The final models for explaining observed haemolymph sugars and lipid contents and their effects are summarised in Table 1. The significance of fixed effects in the final models was assessed with Type II Wald *χ*^*2*^ tests using the *Anova* function of the *car* package [36]. Conditional cR^2^ and marginal mR^2^ were calculated using the *performance* package [37]. Pairwise comparisons between ‘age’ classes and ‘head width’ slopes (slopes only for the absolute lipid content model, which retained an interaction term) were conducted using the *emmeans* package with Holm-Bonferroni corrections.

**Table 1.**
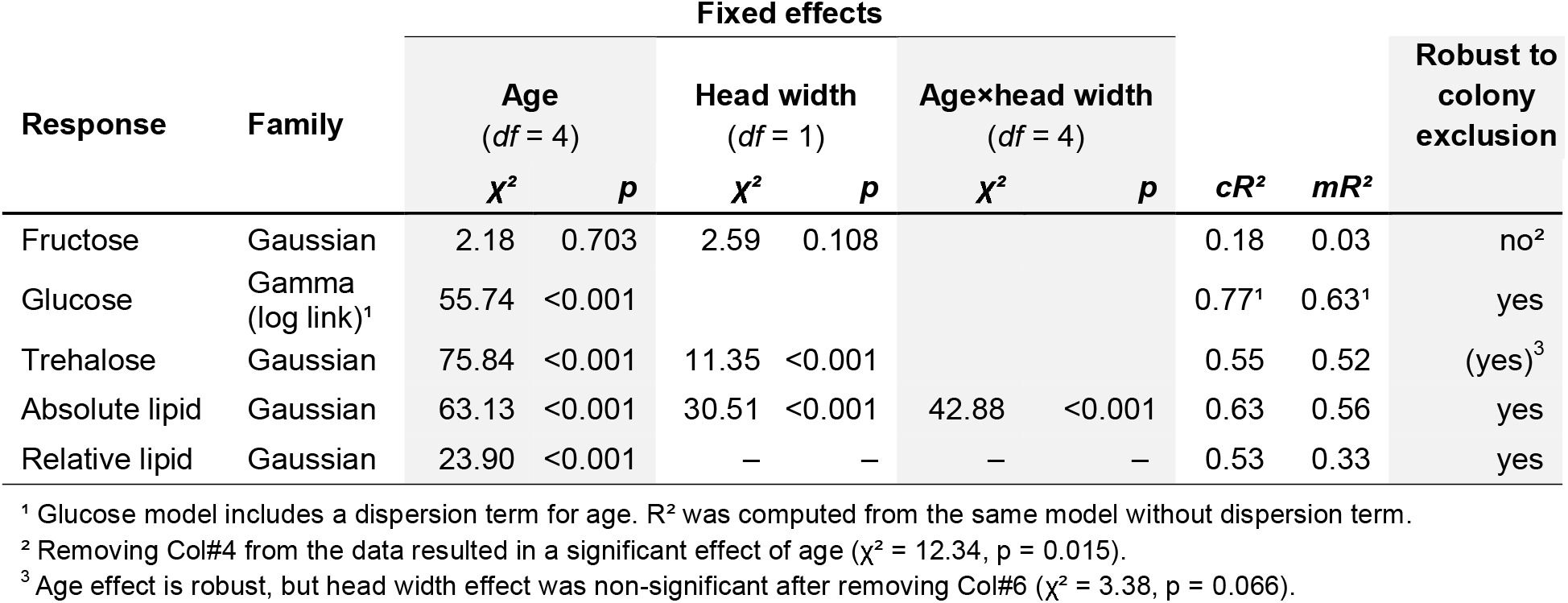
Final generalised linear mixed models (GLMMs) for haemolymph fructose, glucose and trehalose concentrations (μg/μL), absolute (mg) and relative (%) lipid content in bumblebee workers (*n* = 159) acros five age classes. All models included colony and collection date as random intercepts. Effects of age, head width (proxy for body size) and their interaction were tested with Type II Wald *χ*^2^ tests. Conditional (cR^2^) and marginal (mR^2^) R^2^ show the variance explained by fixed plus random effects and by fixed effects alone, respectively. Sensitivity of age and head width effects to colony composition was assessed by refitting each model after excluding one colony at a time (detailed results in Table S2).

|  |  | **Fixed effects** |  |  |  |  |  |  |  |  |
|---|---|---|---|---|---|---|---|---|---|
| **Response** | **Family** | **Age**<br>(*df* = 4) |  | **Head width**<br>(*df* = 1) |  | **Age×head width**<br>(*df* = 4) |  |  |  | **Robust to colony exclusion** |
|  |  | ***χ²*** | ***p*** | ***χ²*** | ***p*** | ***χ²*** | ***p*** | ***cR²*** | ***mR²*** |  |
| Fructose | Gaussian | 2.18 | 0.703 | 2.59 | 0.108 |  |  | 0.18 | 0.03 | no² |
| Glucose | Gamma<br>(log link)¹ | 55.74 | <0.001 |  |  |  |  | 0.77¹ | 0.63¹ | yes |
| Trehalose | Gaussian | 75.84 | <0.001 | 11.35 | <0.001 |  |  | 0.55 | 0.52 | (yes)<sup>3</sup> |
| Absolute lipid | Gaussian | 63.13 | <0.001 | 30.51 | <0.001 | 42.88 | <0.001 | 0.63 | 0.56 | yes |
| Relative lipid | Gaussian | 23.90 | <0.001 | – | – | – | – | 0.53 | 0.33 | yes |

¹ Glucose model includes a dispersion term for age. R² was computed from the same model without dispersion term.
² Removing Col#4 from the data resulted in a significant effect of age (χ² = 12.34, p = 0.015).
<sup>3</sup> Age effect is robust, but head width effect was non-significant after removing Col#6 (χ² = 3.38, p = 0.066).

## 3. Results

### 3.1 Relationships between body size proxies, age and colony

There was a strong positive association between the body size proxies ‘head width’ and ‘dry mass’ (GLMM: *χ*^*2*^ = 241.432, *df* = 1, *p* < 0.0001; Pearson’s correlation: *r* = 0.77, *t* = 14.945, *df* = 157, *p* < 0.0001; figure 1a) and ‘head width’ and ‘ITD’ (*χ*^*2*^ = 711.023, *df* = 1, *p* < 0.0001; Pearson’s correlation: *r* = 0.87, *t* = 22.577, *df* = 157, *p* < 0.0001, figure 1b), which was affected by ‘age’ class (GLMM_DryMass_: *χ*^*2*^ = 18.692, *df* = 4, *p* = 0.0009; interaction age×dry mass: *χ*^*2*^ = 11.121, *df* = 4, *p* = 0.0252; GLMM_ITD_: *χ*^*2*^ = 9.991, *df* = 4, *p* = 0.0406) without any significant difference between age classes (all pairwise comparisons: p > 0.05). However, testing the effect of age class on dry mass alone (*χ*^*2*^ = 12.64, *df* = 4, *p* = 0.013), pairwise comparisons revealed a significant difference in dry mass between callows and 28-day-old workers (*p* = 0.040, all other comparisons *p* > 0.05; supplementary figure S1b). There was no significant difference in head width between colonies (*χ* ^*2*^ = 7.120, *df* = 5, *p* = 0.212; supplementary figure S1a).

### 3.2 Effects on haemolymph sugar concentrations

There was significant effect of worker age on both glucose (*χ*^2^_age_ = 55.740, *df* = 4, *p* < 0.0001; Table 1) and trehalose concentrations (*χ*^2^_age_ = 75.839, *df* = 4, *p* < 0.0001; Table 1). Pairwise comparisons revealed significant differences between callows and older workers of the 7-, 14-, 21-, and 28-day-old cohort for both glucose (*p* < 0.0001; figure 2*b*) and trehalose (*p* < 0.0001; figures 2*c*), while all other comparisons between older workers were non-significant (*p* > 0.05), except for the comparison of trehalose concentrations between 14- and 21-day-old workers (*p* = 0.032; figures 2*c*). Glucose concentrations were on average approximately 6-fold higher (95% CI 3.6-9.3; 6.43 μg/μL) in older workers than in callows, and trehalose concentrations were on average approximately 4-fold higher (32.9 μg/μL, 95% CI 25.0-40.7; supplementary Table S2). While head width had no significant effect on glucose (LRT model comparison with and without ‘head width’: *χ*^2^ = 1.996, *df* = 1, *p* = 0.158), head width affected trehalose concentrations (*χ*^2^ _head width_ = 11.350, *df* = 1, *p* = 0.0008; Table 1) and showed a generally positive relationship (Pearson’s correlation: *r* = 0.38, *t* = 5.159, *df* = 157, *p* < 0.0001; figure 3*c*). The effect of head width on trehalose, however, was not robust to refitting the final model without colony Col#6 (*χ*^2^_head width_ = 3.381, *df* = 1, *p* = 0.066; supplementary Table S2). We observed no significant effect of either age or head width on fructose concentrations (*χ*^2^_age_ = 2.176, *df* = 4, *p* = 0.703; *χ*^2^_head width_ = 2.589, *df* = 1, *p* = 0.108; Table 1, figures 2a and 3a), but when refitting the final model without Col#4 there was a significant effect of age (*χ*^2^_age_ = 12.342, *df* = 4, *p* = 0.015; supplementary Table S2). The age effects on glucose and trehalose were robust to the exclusion of any single colony (all *p* < 0.0001; supplementary Table S2).

**Figure 2.**
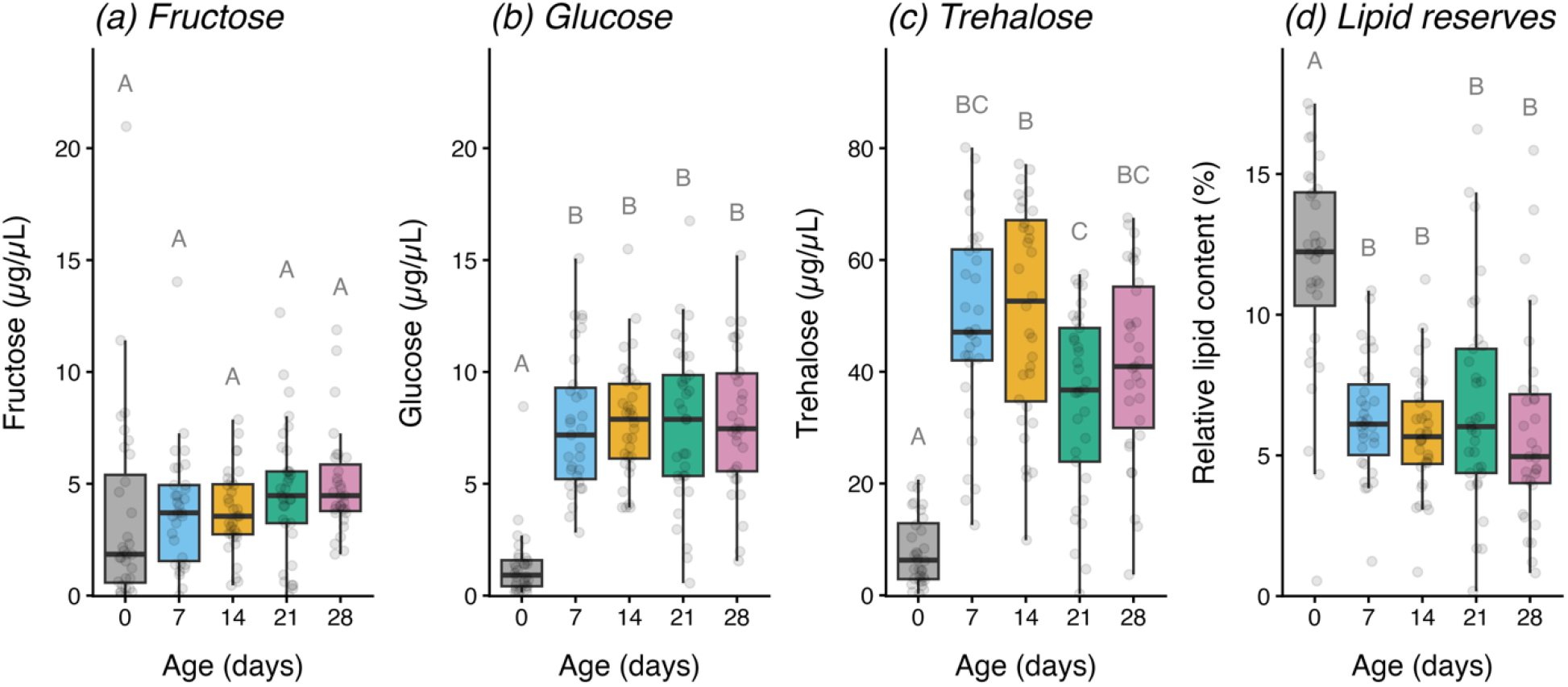
Age-dependent haemolymph sugar levels and relative body lipids in *Bombus terrestris* workers across five age classes. Concentrations (μg/μL) of ***(a)*** fructose, ***(b)*** glucose and ***(c)*** trehalose, and ***(d)*** relative lipid content (% of dry mass) in callows (≤ 1 day old) and 7-, 14-, 21-, 28-day-old workers (*n* = 32, 31, 32, 33 and 31; 4-colonies; for details see supplementary Table S1 and sensitivity assessment Table S2). Absolute lipid content and dry mass across age classes are shown in supplementary figure S1. Boxes show medians and interquartile ranges, and points show individual workers. Different letters indicate significant differences (*p* < 0.05; two-sided pairwise comparisons of estimated marginal means from the GLMMs in Table 1, Holm-adjusted). Groups sharing the same letter do not differ significantly.

**Figure 3.**
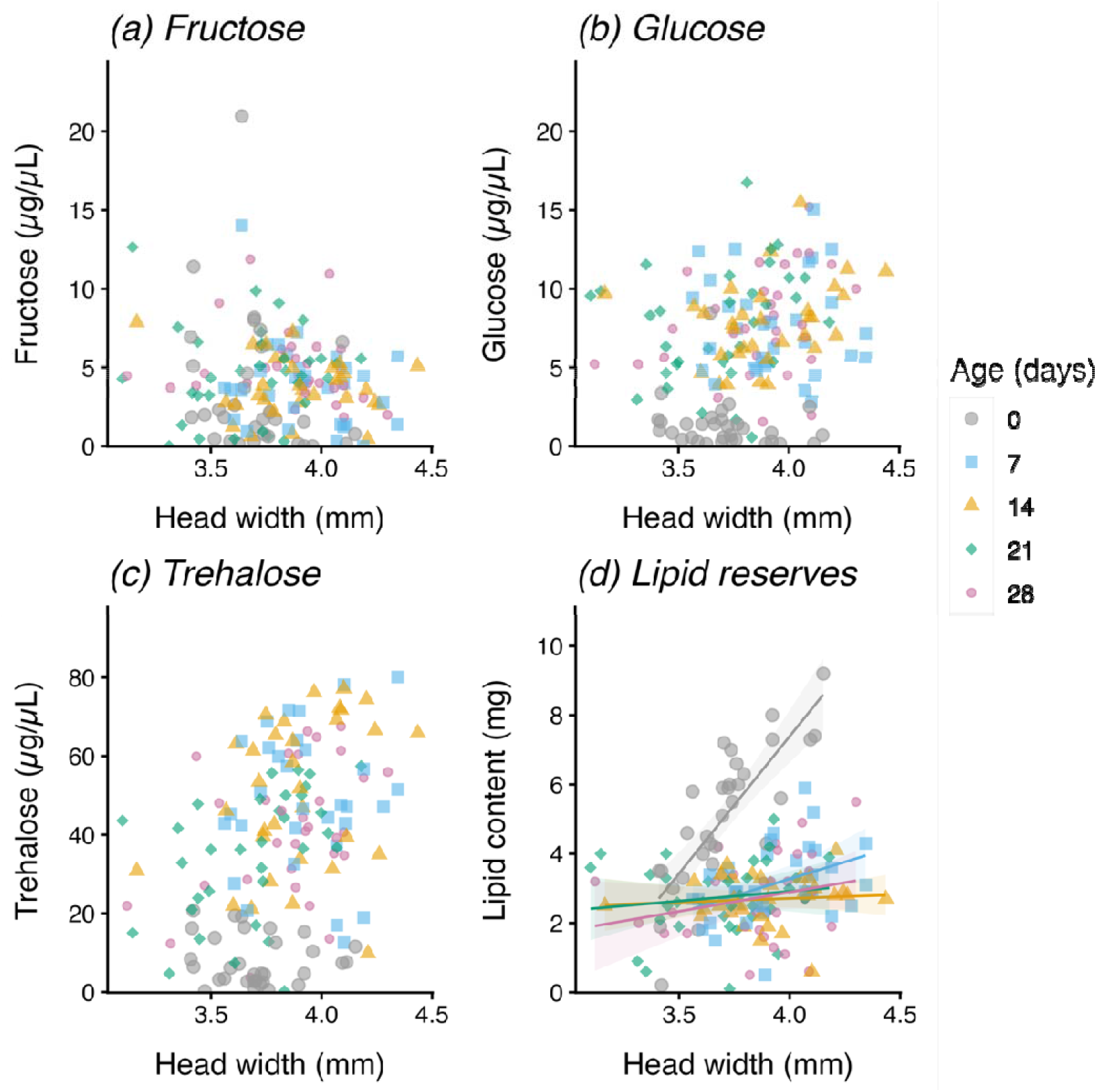
Relationships between head width as proxy for body size and *(a-c)* haemolymph sugar concentrations and *(d)* absolute lipid content in *Bombus terrestris* workers from five age classes. Concentrations (μg/μL) of ***(a)*** fructose, ***(b)*** glucose and ***(c)*** trehalose relative to individual head width (mm). Callows separate clearly from older workers in for glucose and trehalose (figure 2). Head width was not related to fructose (GLMM: *χ*^*2*^ = 2.59, *df* = 1, *p* = 0.11) and was not retained in the final glucose model. Head width was positively related to trehalose (GLMM: *χ*^*2*^ = 11.35, *df* = 1, *p* = 0.0008), but this effect was not robust to the exclusion of Col#6 (*χ*^*2*^ = 3.38, *p* = 0.066; supplementary Table S2). ***(d)*** Absolute lipid content (mg) increased with head width in callows (slope 7.52 mg/mm, 95% CI 5.61–9.42) and weakly in 7-day-old workers (2.28 mg/mm, 95% CI 0.71–3.85), but not in older workers (GLMM: age×head width: *χ*^*2*^ = 42.88, *df* = 4, *p* < 0.001). Age classes are denoted by distinct symbols and colours: callows (≤ 1 d; grey circles, *n* = 32), 7 d (sky blue squares, *n* = 31), 14 d (orange triangles, *n* = 32), 21 d (bluish-green diamonds, *n* = 33), and 28 d (reddish-purple small circles, *n* = 31). Lines in *(d)* show linear regressions with 95% confidence intervals for each age class. Statistics are from the GLMMs in Table 1.

### 3.3 Effects on absolute and relative lipid content

There was a significant effect of worker age (*χ*^2^_age_ = 63.132, *df* = 4, *p* < 0.0001), head width (*χ*^2^_head width_ = 30.513, *df* = 1, *p* < 0.0001), and their interaction on absolute lipid content (*χ*^2^_age×head width_ = 42.879, *df* = 4, *p* < 0.0001; Table 1). Pairwise comparisons revealed significant differences between callows and older workers of the 7-, 14-, 21-, and 28-day-old cohort (*p* < 0.0001;supplementary figure S1*c*), while all other comparisons between older workers were non-significant (*p* > 0.05). While absolute lipid content increased with head width in callows (slope 7.52 mg/mm, 95% CI 5.61–9.42, figure 3d) and weakly in 7-day-old workers (2.28 mg/mm, 95% CI 0.71–3.85), this was not apparent in older workers (14 d: 0.19 mg/mm, 95% CI −1.19 to 1.56; 21d: 0.96 mg/mm, 95% CI −0.46 to 2.39; 28d: 1.15 mg/mm, 95% CI −0.26 to 2.56). Likewise, age significantly affected the relative lipid content (*χ*^2^_age_ = 23.904, *df* = 4, *p* < 0.0001; Table 1, figure 2*d*), with callows having a higher relative lipid content than older workers (difference between older workers and callows: −5.68%, 95% CI −8.16 to −3.20; supplementary Table S2). As the relative lipid content already accounts for bee dry mass, this model did not include head width as an additional body size proxy.

## Discussion

Our study provides empirical evidence of a significant shift in the energy stores of *B. terrestris* between callows (newly emerged workers, ≤ 1 day) and older workers (7-28 days), characterised by an increase in haemolymph glucose and trehalose levels (figure 2*b,c*) alongside a significant reduction of absolute and relative body lipid content (figure 2*d*, supplementary figure S1*c*). Interestingly, haemolymph sugars and lipid contents did not significantly differ between the 7-, 14-, 21-, and 28-day-old workers, with the exception of trehalose, which was significantly lower in 21-day-older worker compared to 14-day-old workers. This transition from a lipid-rich, low-sugar state to a low-lipid, sugar-rich state therefore appears to be largely completed within the first week of adult life. Because we sampled only callows and 7-day-old workers, but not within this period, we cannot say when exactly the transition occurs or whether it is gradual or abrupt.

The approximately 6-fold and 4-fold increase in glucose and trehalose levels, respectively, in older workers (7-28 days) compared to callows (figure 2*b,c*) align with patterns observed in honeybees, where foragers maintain trehalose levels twice as high as those of nurse and newly emerged workers [17]. In honeybees, foraging activity and flight capacity peak between the age of 15 and 32 days, depending on the season [38, 39]. Although this is consistent with the idea that workers must first mobilise circulating energy resources before being able to fly, our haemolymph measurements do not fully support the previously reported lower flight performance in 7-day-old workers compared to 14- and 21-day-old bumblebee workers [7]. Fuel availability might therefore not the limiting factor, and other aspects such as flight muscle maturation may be more important [40]. However, some limitations of our study need to be considered. Feeding all bees with 40% sucrose solution for 3 hours before sampling for standardisation, rather than baseline (fasted) or field levels, may have influenced our haemolymph sugar measurements. As ingested sucrose is hydrolysed to glucose and fructose before it can be converted to trehalose [12], this may have elevated particularly glucose and fructose levels. Because we did not measure food consumption, we cannot account for the possibility that callows consumed less sucrose solution than older workers during this period, and therefore, their lower glucose and trehalose levels may partly reflect lower recent food intake rather than a physiological difference alone. However, there was no difference in fructose levels between callows and older workers, suggesting that the observed effect cannot be explained by food intake alone. There might also be a slight selection bias among older workers, as not all workers will reach the age of 28 days, whereas others naturally live much longer [41], and chronological age may not always reflect physiological age. Furthermore, colonies contributed unequal numbers of workers and callows originated from only four colonies. Although we included colony ID as a random intercept and the age effects on glucose, trehalose and lipid content were robust to the exclusion of any single colony, the absence of an age effect on fructose depended on one colony (Col#4) and should be interpreted with caution (supplementary Table S2). Lastly, haemolymph collection itself may have induced variation between samples, although we carefully collected clean, uncontaminated haemolymph (only one sample with increased sucrose levels of 0.55 μg/μL, which was excluded from our data).

Despite age being the primary explanatory variable, we found a positive relationship between body size (i.e. head width) and trehalose levels (figure *3c*, Table 1). This association was rather weak, inconsistent across age class, and not robust to refitting the model when excluding one colony (Table 1, supplementary Table S2), and therefore, should be regarded as exploratory. If this relationship is real, elevated trehalose levels with increasing body size may be required for higher maximum and mean flight velocities observed in larger workers [7], which we did not measure in the present study. We expected body size to influence haemolymph sugar levels because metabolic rate and energy demand increase with body mass, and approximately 50% of the variability in the flight metabolic rate in bumblebee workers is explained by body size [28]. However, this does not necessarily translate into haemolymph sugar concentrations as previous calculations suggest relatively constant energy demand in bumblebees regardless of flight speed, as increased power demands are offset by a reduction in the power needed to generate lift [42]. More recent research confirms that the metabolic rate does not significantly change across various wind speeds bumblebees faced in tethered or free flight [43]. Therefore, it remains unclear whether larger workers require higher haemolymph sugar concentrations to fuel faster flight. Moreover, haemolymph sugar concentrations reflect the balance between uptake, synthesis and use rather than the pure size of the energy stores. High variability in haemolymph sugar concentrations has been reported among returning foraging bumblebees in the field, likely reflecting environmental factors such as flower availability [29]. Regardless, the large difference between callows and older workers accounted for most of the variance while body size (head width) only played a minor role in our trehalose data.

While haemolymph sugar levels increase substantially in older bumblebee workers, their relative and absolute whole-body lipid reserves were simultaneously reduced by roughly half compared to callows (figure 2*d*, supplementary figure S1*c*). This finding is consistent with the reduction of lipid reserves prior to the transition from nursing to foraging in honeybees [16], a period when lipid synthesis decreases (around the age of 8 days) [44]. It has been suggested that the reduction of lipids, which goes along with a reduction in body mass, is an important adaptation for increasing flight performance in honeybees [38, 39]. Indeed, we found a significant interaction between absolute lipid content and the body size proxy head width as well as a strong positive correlation in callows which disappears as workers age (figure 3*d*). While we found a strong positive linear relationship between body size proxies head width and ITD (figure 1*b*), dry mass, which was used to calculate the relative lipid content, may not scale linearly in callows (figure 1*a*) as shown for the relationship between head width and body mass in newly emerged worker and male bumblebees [41], and therefore, may have influenced our results of the relative lipid contents (figure 2*d*).

Because foraging flights are energetically costly and high activity is known to reduce longevity [4, 25-27, 45], the energetic state may influence the division of labour in bumblebees [6, 22]. It has been suggested that larger workers may avoid energy-intensive tasks as they age to reserve lipid stores for oocyte development [6, 22]. In our data, larger workers did not have higher absolute lipid contents than smaller workers from 14 days of age onwards (figure 3*d*), but because we did not observe their behaviour or assess ovary development, this cannot be interpreted as evidence against such a strategy. Worker reproduction in bumblebees appears to be socially suppressed through pheromonal signalling and physical dominance of the queen during the ergonomic growth phase until the ‘switching point’ is reached, at which mostly unfertilized eggs (male offspring) are laid both by the queen and some workers [6, 8, 9, 22]. Because we did not assess colony stage and ovary development, we do not know whether we sampled workers that were reproductively active. Nonetheless, our data suggest that total body lipid content scales with body size (head width) in callows and that this relationship weakens or disappears in older workers. Whether this reflects an increased resource and energy demand needed for processes such as flight muscle maturation [46], ovary development [11, 47] or cuticular hydrocarbon (CHC) synthesis [48] cannot be resolved with our data.

In conclusion, our findings show a pronounced shift in energy stores within the first week of adult life in *B. terrestris* workers, from high body lipid reserves and low haemolymph sugar levels to lower lipid reserves and high glucose and trehalose levels. Lipid reserves and haemolymph sugar levels were relatively constant across older age classes (7-28 days), and body size proxies (head width, which was highly correlated with ITD and dry mass) explained little additional variation. This transition is consistent with maturation processes that may contribute to task allocation in bumblebees, but because we did not measure behaviour, ovary development, haemolymph protein or hormone titres in the present study, links between these physiological changes, task allocation and worker reproduction remain to be tested. As newly emerged workers are physiologically distinct from older workers, we suggest that age should be routinely considered when conducting experiments in behavioural and evolutionary ecology.

## Data availability

Dataset and R script used for statistical analysis and producing all figures are available from the Zenodo Repository: 10.5281/zenodo.18282075.

## Acknowledgments

We thank Prof. Dr. Joachim Ruther for fruitful discussion that helped us to optimise our derivatisation protocol, Prof. Dr. Tomer J. Czaczkes for sponsoring bumblebee colonies, and Prof. Dr. Reinhard Sterner for allowing us to use the lyophiliser. Furthermore, we thank three anonymous reviewers who have helped us improving the overall quality of our manuscript. We used Claude from Anthropic for proofreading.

## Author contributions

H.S.: investigation, data curation. C.B.: investigation. E.S.: methodology, resources, writing – review and editing. C.K.: conceptualization, methodology, investigation, data curation, validation, formal analysis, writing – original draft preparation, writing – review and editing, visualization, supervision, project administration.

## References

1. Gordon DM. From division of labor to the collective behavior of social insects. Behavioral Ecology and Sociobiology. 2016;70:1101–8.

2. Duarte A, Weissing FJ, Pen I, Keller L. An evolutionary perspective on self-organized division of labor in social insects. Annual Review of Ecology, Evolution, and Systematics. 2011;42:91–110.

3. Johnson BR. Division of labor in honeybees: form, function, and proximate mechanisms. Behavioral Ecology and Sociobiology. 2010;64:305–16.

4. Brian AD. Division of labour and foraging in Bombus agrorum Fabricius. The Journal of Animal Ecology. 1952:21(2):223–40.

5. Goulson D, Peat J, Stout JC, Tucker J, Darvill B, Derwent LC, et al. Can alloethism in workers of the bumblebee, Bombus terrestris, be explained in terms of foraging efficiency? Animal Behaviour. 2002;64(1):123–30.

6. Jandt JM, Dornhaus A. Competition and cooperation: bumblebee spatial organization and division of labor may affect worker reproduction late in life. Behavioral Ecology and Sociobiology. 2011;65:2341–9.

7. Gilgenreiner M, Kurze C. Age dominates flight distance and duration, while body size shapes flight speed in Bombus terrestris L.(Hymenoptera: Apidae). Proceedings of the Royal Society B: Biological Sciences. 2024;291(2027):20241001. doi: 10.1098/rspb.2024.1001.

8. Bourke AFG, Ratnieks FLW. Kin-selected conflict in the bumble-bee Bombus terrestris (Hymenoptera: Apidae). Proceedings of the Royal Society B: Biological Sciences. 2001;268(1465):347–55. doi: 10.1098/rspb.2000.1381.

9. Alaux C, Jaisson P, Hefetz A. Regulation of worker reproduction in bumblebees (Bombus terrestris): workers eavesdrop on a queen signal. Behavioural Ecology and Sociobiology. 2006;60(3):439–46.

10. Zhuang M, Colgan TJ, Guo Y, Zhang Z, Liu F, Xia Z, et al. Unexpected worker mating and colony-founding in a superorganism. Nature Communications. 2023;14(1):5499.

11. Arrese EL, Soulages JL. Insect Fat Body: Energy, Metabolism, and Regulation. Annual Review of Entomology. 2010;55:207–25.

12. Thompson SN. Trehalose □ The insect ‘blood’ sugar. In: Simpson SJ, editor. Adv Insect Physiol, Vol 31. Advances in Insect Physiology. 312003. p. 205–85.

13. Suarez RK. Energy metabolism during insect flight: biochemical design and physiological performance. Physiological and Biochemical Zoology. 2000;73(6):765–71.

14. Robinson GE. Regulation of honey bee age polyethism by juvenile hormone. Behavioral Ecology and Sociobiology. 1987;20:329–38.

15. Schilcher F, Scheiner R. New insight into molecular mechanisms underlying division of labor in honeybees. Current Opinion in Insect Science. 2023; 59:101080.

16. Toth AL, Kantarovich S, Meisel AF, Robinson GE. Nutritional status influences socially regulated foraging ontogeny in honey bees. Journal of Experimental Biology. 2005;208(24):4641–9.

17. Mayack C, Phalen N, Carmichael K, White HK, Hirche F, Wang Y, et al. Appetite is correlated with octopamine and hemolymph sugar levels in forager honeybees. Journal of Comparative Physiology A. 2019;205(4):609–17. PubMed PMID: Mayack2019.

18. Blatt J, Roces F. Haemolymph sugar levels in foraging honeybees (Apis mellifera carnica): dependence on metabolic rate and in vivo measurement of maximal rates of trehalose synthesis. Journal of Experimental Biology. 2001;204:2709–16.

19. Kurze C, Mayack C, Hirche F, Stangl GI, Le Conte Y, Kryger P, et al. Nosema spp. infections cause no energetic stress in tolerant honeybees. Parasitology Research. 2016;115:2381–8. doi: 10.1007/s00436-016-4988-3.

20. Jandt JM, Dornhaus A. Spatial organization and division of labour in the bumblebee Bombus impatiens. Animal Behaviour. 2009;77(3):641–51.

21. O’Donnell S, Reichardt M, Foster R. Individual and colony factors in bumble bee division of labor (Bombus bifarius nearcticus Handl; Hymenoptera, Apidae). Insectes Sociaux. 2000;47:164–70.

22. Cameron SA. Temporal patterns of division of labor among workers in the primitively eusocial bumble bee, Bombus griseocollis (Hymenoptera: Apidae). Ethology. 1989;80(1-4):137–51.

23. Araújo E, Costa M, Chaud-Netto J, Fowler HG. Body size and flight distance in stingless bees (Hymenoptera: Meliponini): inference of flight range and possible ecological implications. Brazilian Journal of Biology. 2004;64:563–8.

24. Kuhn-Neto B, Contrera FA, Castro MS, Nieh JC. Long distance foraging and recruitment by a stingless bee, Melipona mandacaia. Apidologie. 2009;40(4):472–80.

25. Visscher P, Dukas R. Survivorship of foraging honey bees. Insectes Sociaux. 1997;44:1–5.

26. Schmid-Hempel P, Wolf T. Foraging effort and life span of workers in a social insect. The Journal of Animal Ecology. 1988; 57(2):509–21.

27. Rueppell O, Christine S, Mulcrone C, Groves L. Aging without functional senescence in honey bee workers. Current Biology. 2007;17(8):R274–R5.

28. Skandalis DA, Darveau C-A. Morphological and physiological idiosyncrasies lead to interindividual variation in flight metabolic rate in worker bumblebees (Bombus impatiens). Physiological and Biochemical Zoology. 2012;85(6):657–70.

29. Theodorou P, Kühn O, Baltz LM, Wild C, Rasti SL, Bucksch CR, et al. Bumble bee colony health and performance vary widely across the urban ecosystem. Journal of Animal Ecology. 2022;91(10):2135–48. doi: 10.1111/1365-2656.13797.

30. Mayack C, Carmichael K, Phalen N, Khan Z, Hirche F, Stangl GI, et al. Gas chromatography–Mass spectrometry as a preferred method for quantification of insect hemolymph sugars. Journal of Insect Physiology. 2020;127:104115.

31. Laußer S, Kurze C. Impact of thermal stress during pupal development in a key pollinator. Proceedings of the Royal Society B: Biological Sciences. 2025;292(2061):20252029.

32. Wögler L, Kurze C. Experimental short-term heatwaves negatively impact body weight gain and survival during larval development in Bombus terrestris L. (Hymenoptera: Apidae). Biology Open. 2025;14(4). doi: 10.1242/bio.061781.

33. R Core Team. R: A Language and Environment for Statistical Computing. R Foundation for Statistical Computing, Vienna, Austria. URL https://www.R-project.org/; 2025.

34. Brooks ME, Kristensen K, Van Benthem KJ, Magnusson A, Berg CW, Nielsen A, et al. glmmTMB balances speed and flexibility among packages for zero-inflated generalized linear mixed modeling. The R journal. 2017;9(2):378–400.

35. Hartig F. DHARMa: residual diagnostics for hierarchical (multi-level/mixed) regression models. R package https://cran.r-project.org/web/packages/DHARMa/.

36. Fox J, Weisberg S. An R companion to applied regression. Sage; 2018.

37. Lüdecke D, Ben-Shachar MS, Patil I, Waggoner P, Makowski D. performance: An R Package for Assessment, Comparison and Testing of Statistical Models. Journal of Open Source Software. 2021;6(60):3139. doi: 10.21105/joss.03139.

38. Harrison JM. Caste-specific changes in honeybee flight capacity. Physiological Zoology. 1986;59(2):175–87.

39. Vance JT, Williams JB, Elekonich MM, Roberts SP. The effects of age and behavioral development on honey bee (Apis mellifera) flight performance. Journal of Experimental Biology. 2009;212(16):2604–11.

40. Marden JH. Bodybuilding dragonflies: costs and benefits of maximizing flight muscle. Physiological Zoology. 1989;62(2):505–21.

41. Mandlinger MM, Kurze C. Cyclic thermal stress during pupation triggers irreversible carry-over effects in a key pollinator. Communications Biology. 2026;9(1):1228.

42. Dudley R, Ellington CP. Mechanics of forward flight in bumblebees: II. Quasi-steady lift and power requirements. Journal of Experimental Biology. 1990;148(1):53–88.

43. Senior EJ, Tickle P, Walker SM, Askew GN. Energetics of free and tethered flight in bumblebees (Bombus terrestris, Linnaeus 1758). Biology Letters. 2025;21(11). doi: 10.1098/rsbl.2025.0325.

44. Scofield SL, Amdam GV. Fat body lipogenic capacity in honey bee workers is affected by age, social role and dietary protein. Journal of Experimental Biology. 2024;227(18). doi: 10.1242/jeb.247777.

45. Williams JB, Roberts SP, Elekonich MM. Age and natural metabolically-intensive behavior affect oxidative stress and antioxidant mechanisms. Experimental Gerontology. 2008;43(6):538–49.

46. Roberts SP, Elekonich MM. Muscle biochemistry and the ontogeny of flight capacity during behavioral development in the honey bee, Apis mellifera. Journal of Experimental Biology. 2005;208(22):4193–8. doi: 10.1242/jeb.01862.

47. Hoover SER, Higo HA, Winston ML. Worker honey bee ovary development: seasonal variation and the influence of larval and adult nutrition. 2006;176(1):55–63.

48. Blomquist GJ, Ginzel MD. Chemical Ecology, Biochemistry, and Molecular Biology of Insect Hydrocarbons. Annual Review of Entomology. 2021;66:45–60.

