## supplementary for "Age-dependent shift from high lipid content to high haemolymph sugar levels in *Bombus terrestris* (Hymenoptera: Apidae)"

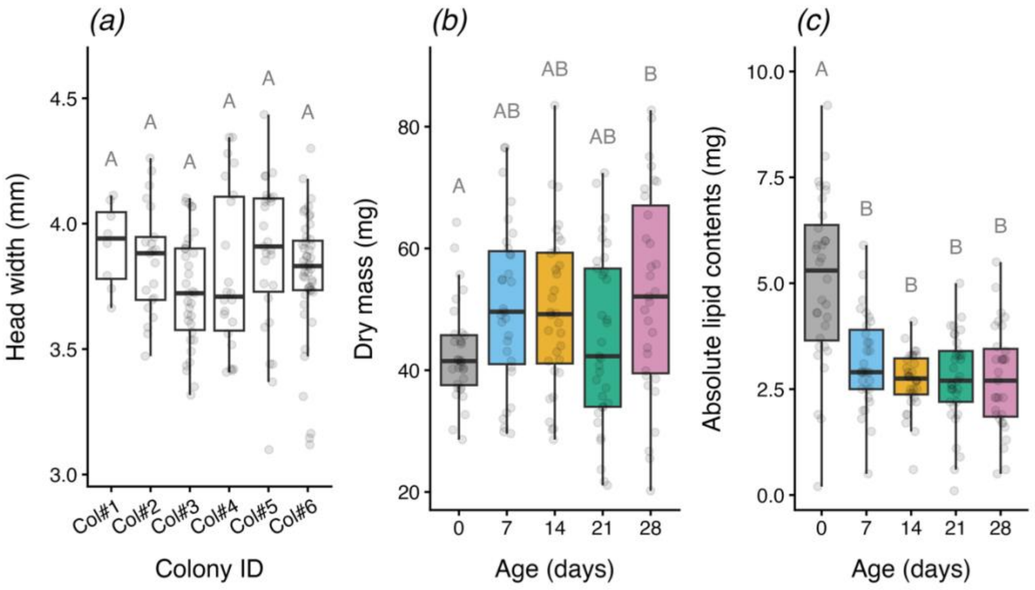


**Figure S1.** Differences in body size and absolute lipid content of *Bombus terrestris* workers between colonies (Col#1–6; *n* = 8, 21, 34, 22, 27 and 47) and age classes (0, 7, 14, 21 and 28 days; *n* = 32, 31, 32, 33 and 31). ***(a)*** Head width (mm), a proxy for body size, did not differ significantly among colonies (linear model, $\chi^2 = 7.12$, $df = 5$, $p = 0.21$). ***(b)*** Dry mass (mg) varied with age (GLMM, $\chi^2 = 12.64$, $df = 4$, $p = 0.013$), but only callows (≤ 1 day old) and 28-day-old workers differed significantly ($p = 0.040$), with callows being lighter. ***(c)*** Absolute lipid content (mg) was higher in callows than in all older age classes (GLMM, age: $\chi^2 = 63.13$, $df = 4$, $p < 0.001$; age×head width: $\chi^2 = 42.88$, $df = 4$, $p < 0.001$; pairwise comparisons evaluated at mean head width). Models in *(b)* and *(c)* included colony and collection date as random intercepts. Boxes show medians and interquartile ranges, and points show individual workers. Different letters indicate significant differences ($p < 0.05$; two-sided pairwise comparisons of estimated marginal means, Holm-adjusted). Groups sharing the same letter do not differ significantly.

**Table S1. Sample size (n) distribution across the five age classes and six colonies.** Model sensitivity to colony composition is shown in Table S2.

| Age (days) | Colony ID | *n* |
|---|---|---|
| 0 | Col#1 | 6 |
| 0 | Col#2 | 12 |
| 0 | Col#3 | 1 |
| 0 | Col#4 | 13 |
| 7 | Col#1 | 1 |
| 7 | Col#2 | 5 |
| 7 | Col#5 | 9 |
| 7 | Col#6 | 5 |
| 7 | Col#3 | 5 |
| 7 | Col#4 | 6 |
| 14 | Col#2 | 2 |
| 14 | Col#5 | 4 |
| 14 | Col#6 | 13 |
| 14 | Col#3 | 11 |
| 14 | Col#4 | 2 |
| 21 | Col#1 | 1 |
| 21 | Col#2 | 1 |
| 21 | Col#5 | 8 |
| 21 | Col#6 | 15 |
| 21 | Col#3 | 8 |
| 28 | Col#2 | 1 |
| 28 | Col#5 | 6 |
| 28 | Col#6 | 14 |
| 28 | Col#3 | 9 |
| 28 | Col#4 | 1 |

**Table S2. Sensitivity of age and head-width effects to colony composition.** Final models (see Table 1) for haemolymph fructose, glucose and trehalose concentrations (µg/µL), absolute body lipid content (mg), and relative body lipid content (%) were refitted after removing one of the six colonies (Col#1-6) at a time. All refitted models retained colony and collection date as random intercepts. Results of the fulls models (no colony removed; "none") are shown in bold. Number of bumblebee workers (*n*) included in each model. $\chi^2$ and *p* values for the effect of age (df = 4), head width (df = 1), and their interaction (df = 4) are from Type II Wald tests. Effects compare callows (≤ 1 day) with the mean of all older workers (7, 14, 21, and 28 days) and are reported with 95% confidence intervals (CI).

| | Colony removed | | Age | | Head width | | Age×Head width | | | 95% CI | | |
|---|---|---|---|---|---|---|---|---|---|---|---|---|
| | | *n* | $\chi^2$ | *p* | $\chi^2$ | *p* | $\chi^2$ | *p* | Effect§ | lower | upper | *p*‡ |
| **Fructose** | **None** | **159** | **2.18** | **0.7034** | **2.59** | **0.1076** | | | **0.64** | **-1.08** | **2.36** | **0.4654** |
| | Col#1 | 151 | 1.95 | 0.7449 | 2.59 | 0.1075 | | | 0.42 | -1.40 | 2.23 | 0.6495 |
| | Col#2 | 138 | 2.69 | 0.6115 | 1.70 | 0.1927 | | | -0.49 | -2.29 | 1.31 | 0.5895 |
| | Col#3 | 125 | 1.81 | 0.7710 | 2.77 | 0.0958 | | | 0.76 | -1.26 | 2.78 | 0.4561 |
| | Col#4 | 137 | 12.34 | 0.0150 | 2.09 | 0.1482 | | | 2.70 | 0.98 | 4.43 | 0.0024 |
| | Col#5 | 132 | 3.09 | 0.5436 | 2.70 | 0.1003 | | | 0.67 | -1.26 | 2.59 | 0.4931 |
| | Col#6 | 112 | 1.01 | 0.9075 | 0.31 | 0.5757 | | | 0.15 | -1.72 | 2.02 | 0.8742 |
| **Glucose** | **None** | **159** | **55.74** | **<0.0001** | | | | | **5.79** | **3.60** | **9.32** | **<0.0001** |
| | Col#1 | 151 | 51.08 | <0.0001 | | | | | 5.57 | 3.42 | 9.08 | <0.0001 |
| | Col#2 | 138 | 39.22 | <0.0001 | | | | | 4.76 | 2.83 | 8.02 | <0.0001 |
| | Col#3 | 125 | 54.60 | <0.0001 | | | | | 7.07 | 4.16 | 12.01 | <0.0001 |
| | Col#4 | 137 | 58.03 | <0.0001 | | | | | 10.04 | 5.48 | 18.38 | <0.0001 |
| | Col#5 | 132 | 55.83 | <0.0001 | | | | | 5.39 | 3.36 | 8.65 | <0.0001 |
| | Col#6 | 112 | 33.27 | <0.0001 | | | | | 5.13 | 2.89 | 9.11 | <0.0001 |
| **Trehalose** | **None** | **159** | **75.84** | **<0.0001** | **11.35** | **0.0008** | | | **32.85** | **25.02** | **40.68** | **<0.0001** |
| | Col#1 | 151 | 64.44 | <0.0001 | 9.99 | 0.0016 | | | 32.63 | 24.05 | 41.20 | <0.0001 |
| | Col#2 | 138 | 79.00 | <0.0001 | 17.89 | <0.0001 | | | 31.60 | 23.94 | 39.25 | <0.0001 |
| | Col#3 | 125 | 66.80 | <0.0001 | 9.85 | 0.0017 | | | 31.86 | 23.73 | 39.98 | <0.0001 |
| | Col#4 | 137 | 64.96 | <0.0001 | 7.79 | 0.0053 | | | 36.33 | 26.93 | 45.74 | <0.0001 |
| | Col#5 | 132 | 59.80 | <0.0001 | 10.53 | 0.0012 | | | 31.35 | 22.63 | 40.07 | <0.0001 |
| | Col#6 | 112 | 48.48 | <0.0001 | 3.38 | 0.0659 | | | 33.98 | 23.80 | 44.17 | <0.0001 |
| **Absolute lipid** | **None** | **159** | **63.13** | **<0.0001** | **30.51** | **<0.0001** | **42.88** | **<0.0001** | **-3.15** | **-3.83** | **-2.48** | **<0.0001** |
| | Col#1 | 151 | 46.46 | <0.0001 | 22.48 | <0.0001 | 41.80 | <0.0001 | -3.44 | -4.23 | -2.66 | <0.0001 |
| | Col#2 | 138 | 54.30 | <0.0001 | 23.31 | <0.0001 | 32.32 | <0.0001 | -2.94 | -3.68 | -2.20 | <0.0001 |
| | Col#3 | 125 | 68.45 | <0.0001 | 25.63 | <0.0001 | 37.34 | <0.0001 | -3.26 | -3.92 | -2.59 | <0.0001 |
| | Col#4 | 137 | 68.97 | <0.0001 | 18.36 | <0.0001 | 24.07 | <0.0001 | -3.22 | -3.96 | -2.48 | <0.0001 |
| | Col#5 | 132 | 40.76 | <0.0001 | 36.65 | <0.0001 | 31.97 | <0.0001 | -3.16 | -4.70 | -1.61 | <0.0001 |
| | Col#6 | 112 | 100.72 | <0.0001 | 29.17 | <0.0001 | 44.19 | <0.0001 | -3.27 | -3.86 | -2.68 | <0.0001 |
| **Relative lipid** | **None** | **159** | **23.90** | **<0.0001** | | | | | **-5.68** | **-8.16** | **-3.20** | **<0.0001** |
| | Col#1 | 151 | 22.35 | 0.0002 | | | | | -5.24 | -7.60 | -2.87 | <0.0001 |
| | Col#2 | 138 | 24.57 | <0.0001 | | | | | -5.57 | -8.04 | -3.11 | <0.0001 |
| | Col#3 | 125 | 19.50 | <0.0001 | | | | | -5.95 | -8.79 | -3.11 | <0.0001 |
| | Col#4 | 137 | 20.84 | 0.0003 | | | | | -6.34 | -9.30 | -3.38 | <0.0001 |
| | Col#5 | 132 | 35.64 | <0.0001 | | | | | -6.35 | -8.49 | -4.20 | <0.0001 |
| | Col#6 | 112 | 35.70 | <0.0001 | | | | | -6.04 | -8.25 | -3.82 | <0.0001 |

§ Mean differences (older workers − callows) in µg/µL, mg (absolute lipid content) or percentage points (relative lipid content). For glucose, ratios of means (older workers/callows) are presented. For absolute lipid content, the effect was evaluated at mean head width because the model includes an interaction between age×head width.

‡ *p-value* (callows vs. older workers)

**Table S3. Retention times (RT) and ion numbers of the target chemical compounds.** Hexachlorobenzene was used as an internal standard (IS).

| Chemical compound | target ions m/z | RT (min) |
|---|---|---|
| Hexachlorobenzene | 283.8 | 25.6 |
| D-Fructose, 1,3,4,5,6-pentakis-O-(trimethylsilyl)-, O-methyloxime | 307.2; 217.1; 73.1 | 29.7 |
| D(-)-Fructose, pentakis(trimethylsilyl) ether, methyloxime (syn) | 307.2; 217.1; 73.1 | 29.9 |
| D-Glucose, 2,3,4,5,6-pentakis-O-(trimethylsilyl)-, O-methyloxime, (1Z) | 319.2; 73.1; 205.1 | 30.2 |
| D-Glucose, 2,3,4,5,6-pentakis-O-(trimethylsilyl)-, O-methyloxime, (1E) | 319.2; 73.1; 205.1 | 30.6 |
| Sucrose, 8TMS derivative | 361.1; 73.1; 217.1 | 43.1 |
| D-(+)-Trehalose, octakis(trimethylsilyl) ether | 361.2; 191.1; 73.1 | 44.6 |

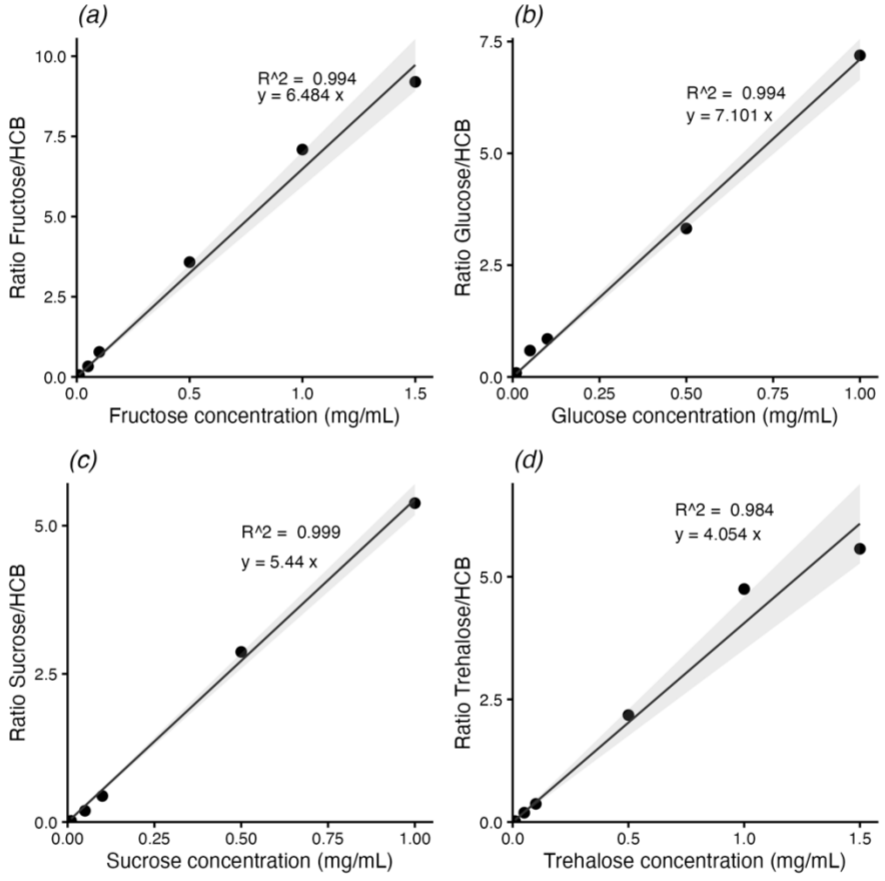


**Figure S2. Standard curves for quantifying sugar levels in haemolymph samples: (*a*) fructose, (*b*) glucose, (*c*) sucrose, and (*d*) trehalose.** Linear regressions confirm highly consistent measurements of the standard sugars relative to Hexachlorobenzene (HCB, used as internal standard) at different concentrations, indicate by $R^2 > 0.95$. Resulting slope formulas were used to calculate concentrations of each sugar in haemolymph samples (for details see methods).
